# Organoid-Based Transcriptomics Indicate IFIT-Associated Immune Modulation during Cryptotanshinone Treatment in Bladder Cancer

**DOI:** 10.64898/2026.01.28.702456

**Authors:** Mengni Yang, Rui Li, Yu Dong, Mengting Zhou, Junning Zhao, Miao Liu, Ruirong Tan

**Author notes:** These authors contributed equally: Mengni Yang, Rui Li, Yu Dong. Correspondence: Ruirong Tan, Miao Liu, Junning Zhao.

## Abstract

Bladder cancer (BC), particularly muscle-invasive and metastatic disease, remains a major clinical challenge despite recent advances in immunotherapy. In this study, we aimed to identify a promising antitumor compound from five candidate small molecules and to explore its potential roles in BC progression. Through antiproliferative screening, cryptotanshinone (CTS) was identified as the promising candidate. Using both two-dimensional BC cell lines and three-dimensional bladder tumor organoid models, we comprehensively evaluated the effects of CTS on cell proliferation, migration, apoptosis, and organoid growth. To further explore the underlying mechanisms, transcriptomic sequencing based on bladder cancer organoid models, protein-protein interaction network analysis, and public databases (TCGA-BLCA, TIMER, and TISIDB) were integrated to examine immune-related pathways and potential molecular targets associated with CTS. GeneMANIA network prediction and molecular docking analyses were subsequently performed to investigate upstream regulatory networks and the potential interactions between CTS and key components of the cGAS-STING-IFN-I-JAK-STAT signaling pathway. Integrative analyses suggested that IFIT1, IFIT2, and IFIT3 may function as immune-associated genes potentially linked to BC progression, patient prognosis, immune cell infiltration, and PD-1/PD-L1 expression. Molecular docking results suggested that CTS may interact with core regulatory proteins within the cGAS-STING-IFN-I-JAK-STAT pathway, potentially influencing IFIT transcriptional regulation. Collectively, these findings indicate that CTS exhibits measurable antitumor and immunomodulatory effects, which may be associated with modulation of the cGAS-STING-IFN-I-JAK-STAT-IFIT signaling axis, supporting its potential as a small-molecule candidate for bladder cancer.

## 1. Introduction

Bladder cancer (BC), the ninth most common cancer globally, typically originates from urothelial cells and is characterized by clinical manifestations such as hematuria and urinary abnormalities[1]; in 2022, there were approximately 614,000 new cases and 220,000 deaths worldwide, with significantly higher incidence and mortality rates in men compared to women[2]. Although progress has been made in understanding its molecular mechanisms, treatment options for muscle-invasive and metastatic bladder cancer remain limited, with current therapies primarily relying on chemotherapy and radical cystectomy[3, 4]. Currently, commonly used chemotherapy regimens, such as cisplatin-based combination therapy, have a response rate of approximately 50-60%. However, relapse rates remain high, severely impacting long-term survival[5]. In recent years, immune checkpoint inhibitors, such as pembrolizumab and nivolumab, have shown promise in the treatment of metastatic bladder cancer, improving survival outcomes in some patients[6, 7]. However, only a subset of patients benefit from these treatments, highlighting the urgent need to develop reliable prognostic biomarkers to better guide treatment decisions. With the increasing understanding of molecular biomarkers, immune checkpoint pathways, and signaling pathway abnormalities, new potential has emerged for the development of prognostic tools and personalized treatment strategies for bladder cancer. Identifying these key biomarkers is crucial for improving treatment efficacy and enhancing patient prognosis.

Organoid technology has emerged as a key tool in cancer research, enhancing our understanding of tumor biology, drug responses, and personalized treatment. Patient-derived bladder cancer organoids faithfully replicate tumor characteristics, the microenvironment, and disease progression, offering a reliable platform for precision medicine[8]. These organoids can model the evolution of tumors from early to late stages and demonstrate how tumor cells respond to treatments like chemotherapy and immunotherapy[9, 10]. This provides valuable insights into tumor dynamics and drug resistance. In immunotherapy research, organoids are used to simulate the tumor-immune microenvironment by incorporating immune cells. This enables more accurate evaluation of therapies like immune checkpoint inhibitors[11]. Organoids also support personalized treatment by predicting patient-specific drug responses, helping to identify the most effective therapies and streamline drug discovery[12]. In conclusion, organoid technology is revolutionizing bladder cancer research, offering new insights for drug development and clinical treatment. Furthermore, our research group has also developed BBN-induced bladder cancer organoid models, providing a valuable platform for studying tumor mechanisms and screening therapeutic candidates.

Cryptotanshinone (CTS) is a lipophilic quinone compound with diverse pharmacological activities, including antitumor and immunomodulatory effects[13]. Previous studies have demonstrated that CTS promotes apoptosis in bladder cancer cells via the PTEN/PI3K/AKT signaling pathway[14] 4 and suppresses tumor cell invasiveness through modulation of the mTOR/β-catenin/N-cadherin axis[15]. Although these findings provide important preliminary insights into the therapeutic potential of CTS in bladder cancer, they are limited by the lack of comprehensive evaluation in experimental models that more accurately recapitulate the biological complexity of this disease. To further investigate the role of CTS in bladder cancer, we employed a BBN-induced bladder cancer organoid model and performed bulk RNA transcriptome sequencing to explore the molecular mechanisms associated with CTS-mediated antitumor effects.

The interferon-induced tetratricopeptide repeat (IFIT) family consists of four members: IFIT1, IFIT2, IFIT3, and IFIT5. Their expression is primarily regulated by interferon-induced JAK-STAT [16, 17] and cGAS-STING[18, 19] signaling pathways. IFITs play crucial roles in antiviral immune responses, cellular function regulation, and tumor initiation and progression[20]. Members of the IFIT family interact with specific molecules through their unique tetratricopeptide repeat (TPR) structure, regulating various biological processes within the cell, including immune responses, cell migration, proliferation, and tumor metastasis[21]. In various cancers, IFIT family members exhibit different roles. For example, overexpression of IFIT3 in pancreatic cancer is closely associated with a “pseudo-inflammatory” phenotype in cells, promoting tumor aggressiveness[22]. Additionally, IFIT3 enhances chemoresistance in cancer cells by regulating post-translational modifications of VDAC2[23]. In pancreatic cancer, IFIT1 promotes cell proliferation, migration, and invasion via the Wnt/β-catenin signaling pathway[24]. In acute myeloid leukemia (AML), the expression levels of IFIT family members are closely related to clinical prognosis, highlighting their biological significance[25]. In skin cutaneous melanoma (SKCM), expression levels of IFIT family members are significantly elevated in tumor tissues[26]. Further bioinformatics analysis suggests that IFIT3 plays a critical role in the tumor immune microenvironment, potentially contributing to immune evasion and metastasis[27]. In conclusion, the IFIT family not only plays an important role in antiviral immune responses but also in immune evasion, tumor microenvironment regulation, and chemoresistance. IFIT family members have potential as biomarkers for cancer and may serve as important therapeutic targets for cancer immunotherapy[28]. IFIT5 promotes EMT, cell migration, and invasion in bladder cancer by downregulating miR-99a and upregulating ICAM1, thereby accelerating tumor metastasis and progression[29]. However, the roles of IFIT1, IFIT2, and IFIT3 in bladder cancer remain underexplored. This study, through transcriptomic sequencing of bladder cancer organoids treated with CTS, identifies IFIT1, IFIT2, and IFIT3 as potential new therapeutic targets for bladder cancer.

In this study, cryptotanshinone (CTS) was selected from five candidate small-molecule compounds based on its inhibitory effects on bladder cancer–associated cellular phenotypes. By integrating a BBN-induced bladder cancer organoid model, bladder cancer cell lines, and bulk RNA sequencing data, we delineated the potential molecular targets of CTS and the biological processes underlying its activity in bladder cancer. Leveraging multidimensional bioinformatics analyses, this study further characterized immune-related signaling programs associated with CTS exposure and generated mechanistic insights that may be informative for future investigations into immunotherapy in bladder cancer.

## 2. Materials and methods

### 2.1. small-molecule compounds

Baicalein, Puerarin, Betulinic acid, Tanshinone IIA, and Cryptotanshinone were purchased from Aladdin (China).

### 2.2. Cell cytotoxicity assay

Cell viability was determined using the Cell Counting Kit-8 (CCK-8, Yeasen Biotechnology, China) in accordance with the manufacturer’s instructions. Initially, 5 × 10³ cells were seeded per well in 100 μL of medium in 96-well plates (Corning, USA). The cells were then exposed to various concentrations of Baicalein, Puerarin, Betulinic acid, Tanshinone IIA, and Cryptotanshinone. After a 24-hour incubation period, 10 μL of CCK-8 reagent was added to each well, followed by an additional incubation of 2 to 4 hours. Absorbance was measured at 450 nm using a microplate reader, with medium-only wells serving as blanks. The absorbance readings were used to represent cell proliferation.

### 2.3. Colony formation assay

1,000 cells of 5637, T24, and UMUC-3 cell lines were seeded into 6-well culture plates and treated with various concentrations of cryptotanshinone (0-4 μM) after 48 hours. The cells were then incubated, with medium changes every 3 days, for a total of 9 days. At the end of the incubation period, the cells were washed with phosphate-buffered saline (PBS). The colonies were subsequently fixed with methanol, stained with a 0.1% crystal violet solution, and analyzed using ImageJ software.

### 2.4. Wound healing assay

5 × 10⁵ 5637 cells were seeded into 6-well plates and cultured until they reached 80–90% confluence. A 200 μL pipette tip was used to create a scratch wound along the midline of each well. The migration of the cells was monitored and photographed using a light microscope (Leica DM IL). The wound area was measured before and after treatment with CTS (0–20 μM) using ImageJ software. The migration rate was expressed as a percentage of the control and was calculated as the proportion of the mean distance between the wound borders relative to the distance that remained cell-free after cell migration.

### 2.5. Flow cytometry

For apoptosis analysis of bladder cancer cells, 5 × 10⁵ 5637 or T24 cells were seeded into 10 cm culture plates and treated with various concentrations of cryptotanshinone (0–20 μM) for 48 hours. After treatment, both attached and floating cells were collected and washed twice with cold PBS. The harvested cells were then stained with Annexin V-PE/7-AAD Apoptosis Detection Kit according to the manufacturer’s instructions and analyzed using flow cytometry.

### 2.6. Organoids construction and cultivation

To generate bladder cancer organoids (BTO) from BBN-induced mouse bladder tissues, begin by digesting the bladder tissues with a collagenase and Y27632 solution to create a single-cell suspension. Afterward, mix the cells with extracellular Matrigel at a 1:2 ratio, aiming for a cell density of 20,000-30,000 cells per drop. Dispense the mixture into culture plates and add an appropriate volume of organoid-specific culture medium. Incubate the plates at 37°C with 5% CO_2_. The medium should be replaced every 3-4 days, and organoids should be passaged every 7-9 days.

### 2.7. Organoid drug sensitivity testing

To seed organoids in a 96-well plate, add 3 μL of cell suspension and 6 μL of Matrigel per well, ensuring a cell density of 450-600 cells per well. Treat the organoids with drug-containing culture medium, using 3 replicates per treatment group, and incubate for 48 hours. On day 3, refresh the medium with fresh drug-containing solution and capture images using a bright-field microscope (Leica). The medium should be replaced again on day 6, followed by another round of imaging. Finally, on day 9, assess organoid viability using the CCK-8 assay (Yeasen, China).

### 2.8. Organoid immunofluorescence

Once the drug-treated organoids are collected, fix them in 4% paraformaldehyde for 45 minutes, followed by permeabilization with 0.3% Triton X-100 for 20-30 minutes. To block nonspecific binding, incubate the organoids in a solution of Normal Goat Serum and BSA at room temperature for 1 hour. Next, apply the primary antibody and incubate overnight at 4°C according to the manufacturer’s instructions. The following day, remove the primary antibody and incubate the organoids with the secondary antibody at room temperature for 1 hour. Finally, use a laser scanning confocal microscope (Zeiss LSM980 with Airyscan2) and confocal dishes (Biosharp) to capture images.

### 2.9. Bulk RNA-sequencing

Bladder cancer organoids treated with CTS or vehicle for 6 days were collected, with three biological replicates in each group. RNA extraction from the samples was carried out using TRI reagent, and the RNA concentration, integrity, and purity were evaluated with the Agilent 5400 Bioanalyzer. Once quality control requirements were met, libraries were constructed and subjected to additional quality assessments prior to sequencing. Subsequent analyses were performed in R (version 4.3.1)[30] using RStudio as the development environment.

### 2.10. PCA analysis

PCA (Principal Component Analysis) is a statistical technique used to simplify complex data by identifying principal features[31]. After filtering raw transcriptome data, sequence alignment was performed with HISAT2 (v2.0.5), and gene expression levels were calculated using featureCounts (1.5.0-p3). Expression profiles in FPKM format were analyzed using the PCA function from the FactoMineR package in R. Visualization was done with ggplot2 and its extensions ggrepel and plyr.

### 2.11. Differential Expression Gene analysis

DEGs (Differentially Expressed Genes) were determined with DESeq2 software (v1.20.0)[32] by applying the thresholds of *p*adj ≤ 0.05 and |log2(fold change)| ≥ 1. Heatmaps and volcano plots were produced using the R packages ggplot2 and pheatmap.

### 2.12. GO and KEGG enrichment analysis

The significant functions and pathways of differentially expressed genes (DEGs) were explored based on gene annotation databases. GO (Gene Ontology)[33] categorizes genes into biological processes, molecular functions, and cellular components, while KEGG[34] (Kyoto Encyclopedia of Genes and Genomes) maps genes to specific pathways. GO functional and KEGG pathway enrichment analyses and visualizations were performed using the R packages clusterProfiler[35], enrichplot, and ggplot2. A p-value less than 0.05 was considered significant for enrichment analysis.

### 2.13. Protein-Protein Interaction (PPI) Network Construction and Analysis

The PPI network was constructed using the STRING database (https://cn.string-db.org) with a confidence score of 0.7, excluding isolated proteins and keeping other parameters at their default settings. The network was subsequently analyzed using Cytoscape (v3.10.2), with further evaluation performed through the cytoNCA plugin[36] to assess the degree centrality (DC), betweenness centrality (BC), and closeness centrality (CC). Core modules in the PPI network were identified using the MCODE plugin[37] (degree cutoff = 2, node score cutoff = 0.2, k-core = 2, max depth = 100). The top three modules contained 16 nodes and 115 edges (score = 15.33), 9 nodes and 16 edges (score = 4), and 4 nodes and 6 edges (score = 4), respectively. Furthermore, we also utilized the GeneMANIA database (http://genemania.org/) to construct a gene network for the *IFIT1/2/3* and to predict potential target genes.

### 2.14. Data Acquisition and Processing for Bioinformatics Analysis

Gene expression data for pan-cancer and adjacent normal tissues, processed into FPKM format using the Toil pipeline[38], were retrieved from The Cancer Genome Atlas (TCGA) and Genotype-Tissue Expression (GTEx) databases through the UCSC XENA platform[39]. Additionally, the FPKM format gene expression data and corresponding clinical information for bladder cancer (BLCA) samples were downloaded and curated from the TCGA database using the R package “TCGAbiolinks”[40].

### 2.15. Pan-Cancer Analyses of IFIT1/2/3 Expression

Data from the TCGA and GTEx databases were merged to compare the mRNA expression levels of *IFIT1/2/3* between tumors and normal tissues across 33 cancer types. Differential expression analysis and visualization of tumors versus normal tissues were performed using the R packages ‘ggplot2’, ‘stats’, and ‘car’. The expression of *IFIT1/2/3* in various cancer cell lines, including different bladder cancer cell lines, was analyzed using the Cancer Cell Line Encyclopedia (CCLE) database[41].

#### Clinical Significance Analysis of IFIT1/2/3 Expression in TCGA-BLCA

Based on TCGA-BLCA expression and clinical data, the expression patterns of *IFIT1/2/3* across bladder cancer subtypes and histologic grades were analyzed. Differential expression analysis and visualization between subgroups were performed using the R packages ‘ggplot2’, ‘stats’, and ‘car’.

### 2.16. Analysis of the Immunological Roles of IFIT1/2/3

ESTMATE: The ESTIMATE algorithm[42] was used to calculate the immune, stromal, and ESTIMATE scores for *IFIT1/2/3* in TCGA-BLCA. BLCA patients were then divided into high and low *IFIT1/2/3* expression groups, and the differences in scores between the two groups were analyzed.

ssGSEA immune infiltration analysis: We applied the ssGSEA algorithm from the GSVA R package (version 1.46.0)[43] to assess immune cell infiltration in TCGA-BLCA, using markers for 24 immune cell types as described in the Immunity article[44]. Additionally, Spearman correlation analysis was performed to examine the association between molecular expression levels and immune cell infiltration. Furthermore, we categorized the BLCA patients into high and low *IFIT1/2/3* expression groups and analyzed the differences in immune cell infiltration levels between the two groups.

TIMER: The Tumor Immune Estimation Resource (TIMER) is a comprehensive tool for analyzing immune infiltration across different cancer types. TIMER 2.0[45], the latest version (http://timer.cistrome.org/), employs six advanced algorithms to provide more reliable estimates of immune infiltration levels in the Cancer Genome Atlas (TCGA). It also offers in-depth analysis and visualization of tumor-infiltrating immune cells. Moreover, the database evaluates the relationship between gene expression and tumor purity. Using TIMER 2.0, we assessed the correlation of IFIT1/2/3 with immune cell infiltration and tumor purity in bladder cancer.

TISIDB: Tumor-Immune System Interaction Database (TISIDB) is a network platform that integrates various types of heterogeneous data to study the interactions between tumors and the immune system. It incorporates data from PubMed, high-throughput screening, exome and RNA sequencing datasets, TCGA, Uniport, GO, DrugBank, and others[46]. In this study, TISIDB was used to investigate the correlation between IFIT1/2/3 and 28 tumor-infiltrating lymphocytes (TILs).

Correlation Analysis: Based on TCGA-BLCA expression data, Spearman correlation analysis was performed to evaluate the relationship between *IFIT1/2/3* and immune-related genes, including chemokines and their receptors, immune inhibitory genes, immune activation genes, MHC molecules and related genes of *IFIT1/2/3*.

### 2.17. Molecular docking

The three-dimensional crystal structure of the protein receptor is primarily obtained from the PDB database (https://www.rcsb.org/), which includes experimentally determined structures. If unavailable, predicted structures from the AlphaFold Protein Structure Database[47, 48], linked to UniProt[49] entries, can be used (Table S1). The protein structure is processed with PyMOL (version 3.1.1) to remove redundant conformations, ligands, and water molecules, and saved in PDB format. It is then prepared for docking using AutoDockTools[50] and converted to PDBQT format.The two-dimensional structure of CTS is retrieved from PubChem (SDF format), converted to MOL2 format using Chem3D, and further processed in AutoDockTools to generate a PDBQT file for docking. Finally, molecular docking of CTS with the related protein is performed using AutoDock Vina[51], and the results are visualized in PyMOL.

### 2.18. Statistical Analysis

All bioinformatics analyses were performed using R v4.3.1. Comparisons between groups were made using the Wilcoxon rank sum test. Survival analyses were conducted using Kaplan-Meier curves and Cox proportional hazards regression models. Spearman correlation analysis was used to evaluate relationships between variables. All experimental statistical analyses were conducted using GraphPad Prism 10.1.1. Differences between two groups were assessed using t-tests, while group comparisons were evaluated using one-way ANOVA.

## 3. Results

### 3.1. CTS exhibits pronounced in vitro anti–bladder cancer activity

To identify candidate compounds with anti–bladder cancer potential, this study evaluated five small-molecule compounds (Fig. 1A), including baicalein (BCA)[52], puerarin (PUE)[53], betulinic acid (BA)[54], tanshinone IIA (Tan-IIA)[55], and cryptotanshinone (CTS)[14], all of which have been reported to exhibit anti-bladder cancer activity — on bladder cancer cell lines 5637 and T24 in vitro. Among these, CTS exhibited the most potent anti-cancer activity, significantly outperforming the other compounds (Fig. 1B-C).

**Fig. 1.**
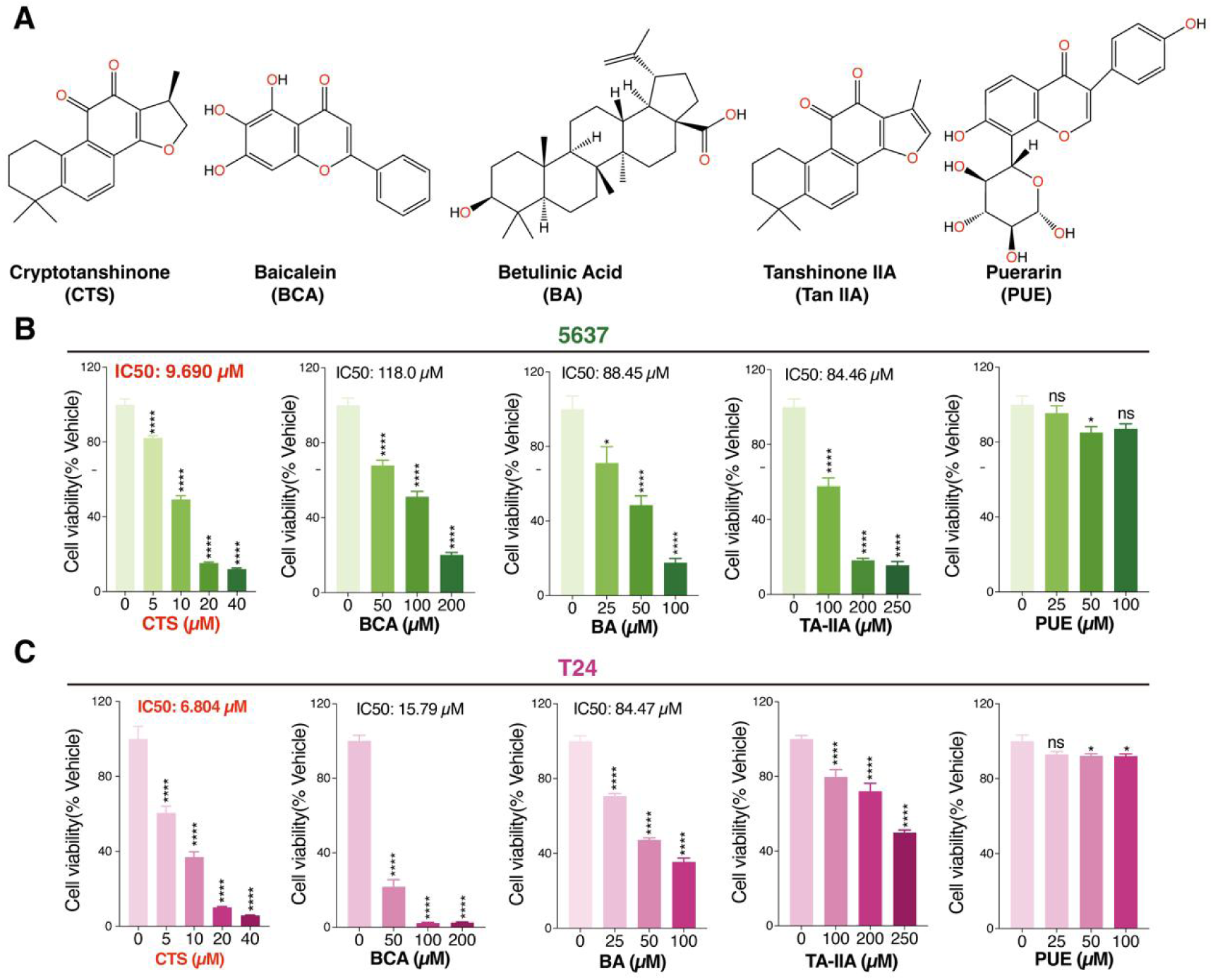
CTS shows comparatively stronger anti–bladder cancer activity in vitro. (A) The chemical structures of Cryptotanshinone (CTS), Baicalein (BCA), Betulinic acid (BA), Tanshinone IIA (Tan-IIA), and Puerarin (PUE). (B) 5637 cells were treated with the indicated concentrations of Cryptotanshinone (CTS), Baicalein (BCA), Betulinic acid (BA), Tanshinone IIA (Tan-IIA), or Puerarin (PUE) for 24 h. Cell viability was determined with the CCK-8 reagent. Data are shown as the mean ± SD, n=6. (C) T24 cells was treated with the indicated concentrations of Cryptotanshinone (CTS), Baicalein (BCA), Betulinic acid (BA), Tanshinone IIA (Tan-IIA), or Puerarin (PUE) for 24 h. Cell viability was determined with the CCK-8 reagent. Data are shown as the mean ± SD, n=6. * *P* < 0.05, ** *P* < 0.01, *** *P* < 0.001, **** *P* < 0.0001. ns, no significance, * compare to Vehicle.

### 3.2. CTS suppresses bladder cancer growth in vitro

Based on the above screening results, subsequent experiments focused on CTS. We first evaluated its antitumor effects using conventional two-dimensional bladder cancer cell models. CTS treatment resulted in a time- and dose-dependent reduction in bladder cancer cell viability (Fig. 2A). Consistently, colony formation assays demonstrated a significant decrease in clonogenic capacity following CTS treatment (Fig. 2B). In addition, wound healing assays showed a dose-dependent inhibition of cell migration upon CTS exposure (Fig. 2C). Apoptosis assays further revealed an increased proportion of apoptotic cells in response to CTS treatment (Fig. 2D). To assess these effects in a more physiologically relevant system, CTS was subsequently examined using BBN-induced bladder tumor organoids (BTOs). Compared with the control group, CTS treatment led to a dose-dependent reduction in both organoid size and number (Fig. 2E–F). Consistently, CCK8 assays demonstrated a significant decline in organoid viability following CTS exposure (Fig. 2G), with an IC₅ ₀ value of 17.3 μM (Fig. 2H). In addition, immunostaining showed reduced expression of the proliferation marker Ki67 in CTS-treated organoids (Fig. 2I). Collectively, these results indicate that CTS exerts inhibitory effects on bladder cancer cell proliferation, migration, and survival across both two-dimensional cell lines and three-dimensional organoid models, supporting its further investigation as a potential therapeutic compound for bladder cancer.

**Fig. 2.**
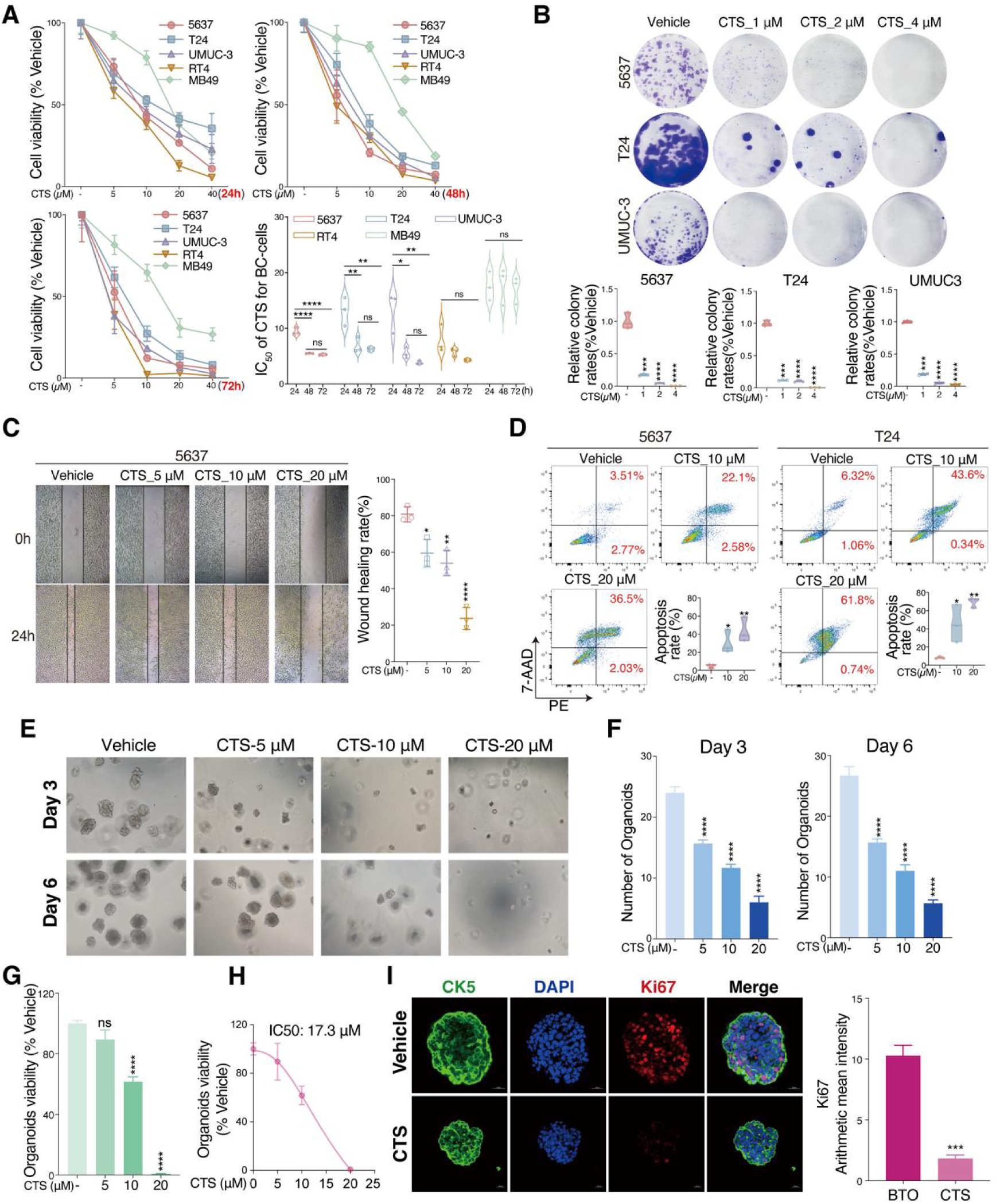
CTS inhibits bladder cancer growth *in vitro*. (A) 5637, T24, RT4, UMUC-3, and MB49 cells were treated with CTS for 24, 48, and 72 hours to determine IC50 values. Cell viability was measured by CCK-8 (mean ± SD, n=6). (B) Anti-proliferation effect of CTS on 5637, T24, and UMUC-3 cells was assessed by colony formation. (C) Wound healing assay of 5637 cells treated with CTS for 24 hours. (D) Apoptosis rates in 5637 and T24 cells after 48h CTS treatment were measured by AnnexinV/7-AAD staining and flow cytometry (mean ± SD, n=3). (E) Bright field micrographs of bladder cancer organoids on Day 3 and Day 6 show that CTS inhibits the growth of bladder cancer organoids in a dose-dependent manner. (F) Day 3 (left) and Day 6 (right), the number of organoids under the same magnification at different CTS doses (mean ± SD, n=3). (G) CCK-8 assay to assess the effect of different doses of CTS on organoid cell viability (mean ± SD, n=6). (H) The IC50 of CTS for BTO. (I) The expression levels of Ki67 in BBN-induced bladder cancer organoids treated with CTS were determined by immunofluorescence staining. * *P* < 0.05, ** *P* < 0.01, *** *P* < 0.001, **** *P* < 0.0001. ns, no significance, * compare to Vehicle. BTO, Bladder Tumor Organoid. CTS, BBN-induced bladder tumor organoids (BTO) treated with CTS.

### 3.3. CTS alters the transcriptomic profile of bladder cancer organoids

To explore the molecular changes associated with CTS treatment in bladder cancer, RNA sequencing (RNA-seq) was performed to compare BBN-induced bladder tumor organoids (BTO) with CTS-treated BTOs (CTS) (Fig. 3A). Principal component analysis (PCA) of the transcriptomic data showed a clear separation between the BTO and CTS groups, indicating distinct global gene expression patterns following CTS treatment (Fig. 3B). Differential expression analysis identified a total of 700 differentially expressed genes (DEGs) between the CTS and BTO groups, including 498 downregulated genes (blue dots) and 202 upregulated genes (red dots) in the CTS group (Fig. 3C). Notably, downregulated genes constituted the majority of DEGs. Heatmap visualization further illustrated consistent expression changes across samples (Fig. 3D), with several genes, including Ifit1, Ifit2, Ifit3, Cxcl15, and Ifi44, showing reduced expression in CTS-treated organoids. These genes have been previously reported to be associated with tumor-related processes such as cell proliferation, migration, and immune regulation. Together, these transcriptomic changes suggest that CTS treatment is accompanied by broad alterations in gene expression programs in bladder cancer organoids, potentially involving pathways relevant to tumor progression and immune-related processes.

**Fig. 3.**
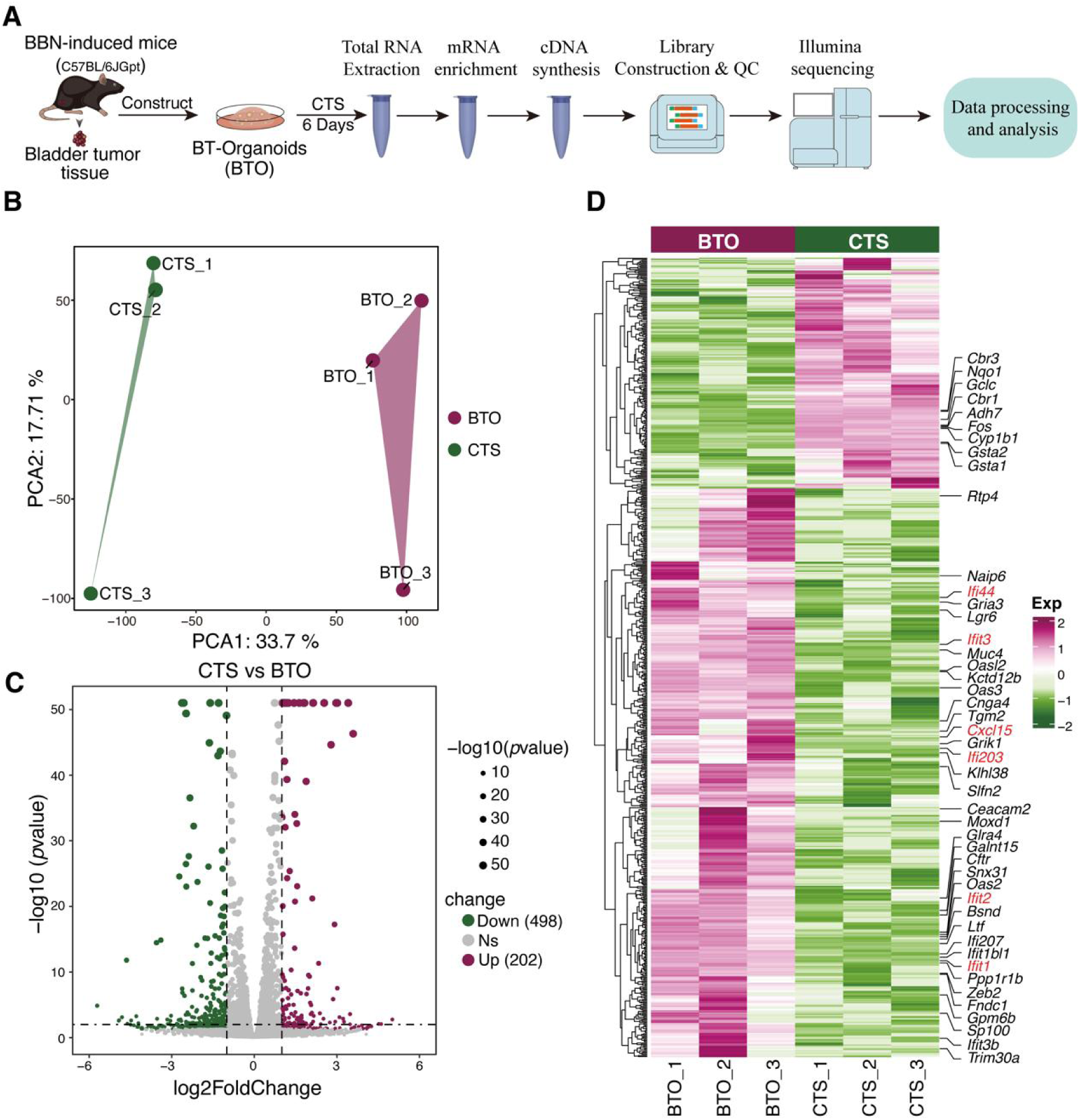
Transcriptomic changes in BTOs following CTS treatment. (A) The workflow of RNA-sequencing. BTO vs CTS, n=3. (B) The PCA of BTO and CTS groups. (C) The volcano plot of DEGs in CTS vs BTO. (D) Heatmap illustrating the DEGs of CTS vs BTO. PCA, Principal Component Analysis. DEGs, Differentially Expressed Genes.

### 3.4. CTS modulates immune-related signaling pathways in bladder cancer organoids

To investigate the biological pathways influenced by CTS treatment in bladder cancer, Gene Ontology (GO) and KEGG pathway enrichment analyses were performed based on the identified differentially expressed genes (DEGs). Upregulated genes were enriched in 170 biological process (BP) terms, 8 cellular component (CC) terms, and 22 molecular function (MF) terms, whereas downregulated genes were enriched in 226 BP terms, 15 CC terms, and 23 MF terms, with downregulated genes showing a broader range of enrichment (Fig. 4A). Functional annotation revealed that upregulated genes were mainly involved in cellular stress responses, detoxification, and metabolic processes. Representative enriched terms included lipid catabolic process, small molecule catabolic process, response to oxidative stress, detoxification, astrocyte projection, L-type voltage-gated calcium channel complex, and oxidoreductase activity (Fig. 4B). In contrast, downregulated genes were predominantly enriched in immune-related processes, including regulation of innate immune response, cytokine-mediated signaling pathway, cellular response to interferon-beta, pattern recognition receptor activity, and CXCR chemokine receptor binding (Fig. 4C). KEGG pathway analysis further supported these observations. Upregulated genes were enriched in 17 pathways (Fig. 4D), primarily related to oxidative stress and metabolic regulation, such as glutathione metabolism, metabolism of xenobiotics by cytochrome P450, drug metabolism–cytochrome P450, and arachidonic acid metabolism (Fig. 4E). Conversely, 13 pathways enriched among downregulated genes (Fig. 4D) were largely linked to immune regulation and tumor-related signaling, including the NOD-like receptor signaling pathway, viral protein interaction with cytokines and cytokine receptors, cAMP signaling pathway, and complement and coagulation cascades (Fig. 4F). Collectively, these enrichment analyses indicate that CTS treatment is accompanied by coordinated alterations in metabolic and immune-related pathways in bladder cancer organoids, with a prominent representation of immune-associated processes among the downregulated gene sets.

**Fig. 4.**
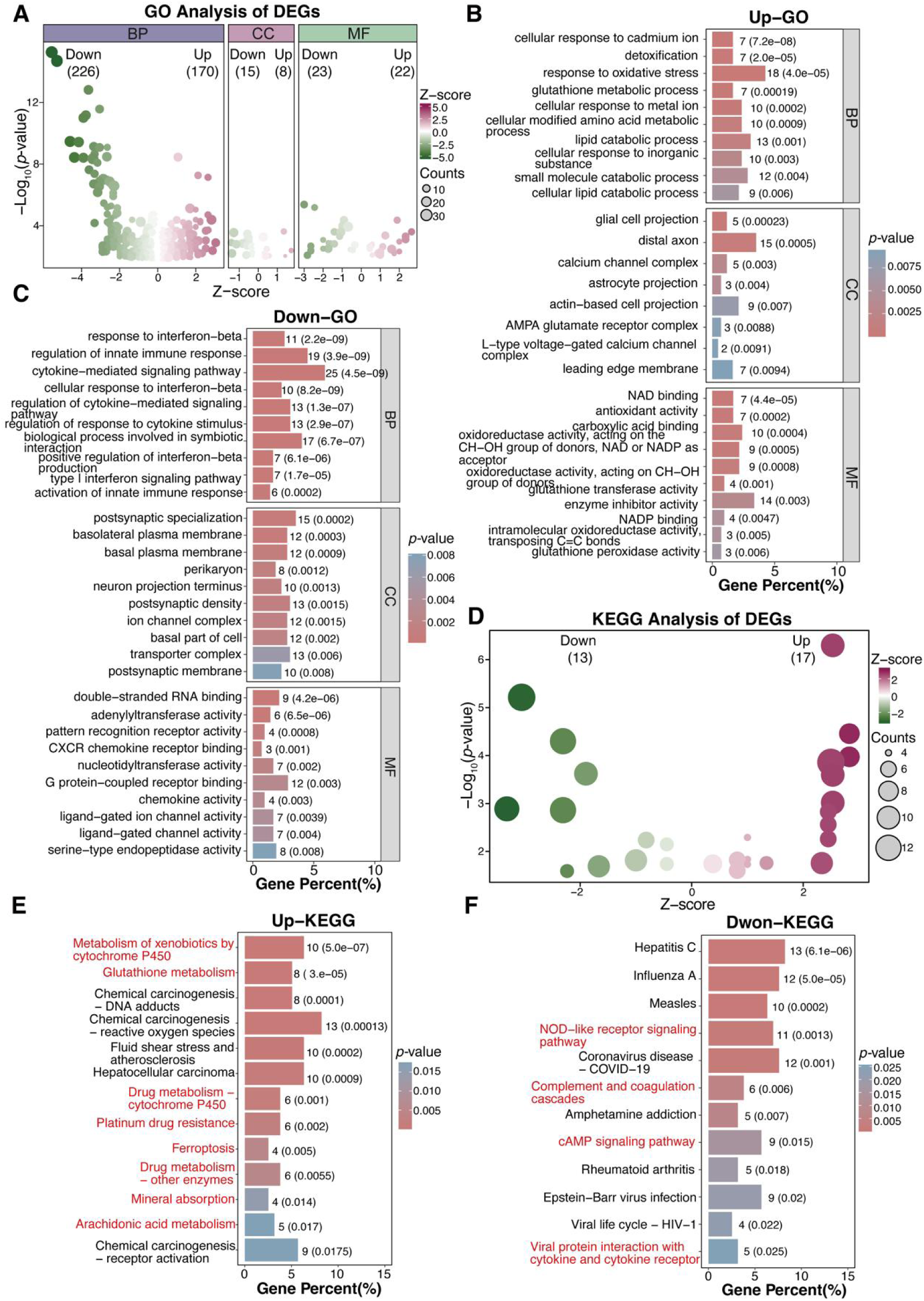
GO function and KEGG pathway enrichment analysis of DEGs. (A) The bubble plot shows the number of GO terms enriched with differentially expressed genes (DEGs), including Biological Process (BP), Cellular Component (CC), and Molecular Function (MF). Z-score < 0 indicates terms enriched with downregulated genes, while Z-score > 0 indicates terms enriched with upregulated genes. (B) GO function enrichment analysis of up-regulated DEGs. (C) GO function enrichment analysis of down-regulated DEGs. (D) The bubble plot shows the number of KEGG pathways enriched with differentially expressed genes (DEGs). Z-score < 0 indicates terms enriched with down-regulated genes, while Z-score > 0 indicates terms enriched with up-regulated genes. (E) KEGG pathway analysis of up-regulated DEGs. (F) KEGG pathway analysis of down-regulated DEGs. GO, Gene Ontology. KEGG, Kyoto Encyclopedia of Genes and Genomes.

### 3.5. The IFIT1/2/3 may represent candidate targets of CTS in bladder cancer

To further explore molecular features potentially involved in CTS responses in bladder cancer, a protein–protein interaction (PPI) network analysis was performed on the differentially expressed genes using the STRING database (confidence score ≥ 0.7). resulting PPI network comprised 120 nodes, including 50 upregulated and 70 downregulated genes, and 296 edges (Fig. 5A). Using the MCODE algorithm, the top three modules within the PPI network were identified, with the highest-ranked module defined as the core module (Fig. 5B-D). This core module included 16 nodes and 115 edges, including genes such as *Ifit3*, *Ifit1*, *Ifi44*, *spP18*, *Rtp4*, *Irf7*, *Isg15*, and *Ifit2* (Table S2). Following CTS intervention, the expression of the top 10 genes in this module was reduced in bladder cancer organoids (Fig. 5E). To assess the clinical relevance of these findings, expression patterns of the corresponding human homologs were examined in the TCGA-BLCA cohort. Notably, these genes were generally expressed at higher levels in tumor tissues compared to adjacent normal tissues. All top 10 genes, except for *IFIT1*, exhibited statistically significant differences (p < 0.001). Although *IFIT1* did not reach statistical significance, it still showed a trend of higher expression in tumor tissues (Fig. 5F). Survival analysis further indicated that higher expression of *IFIT1*, *IFIT2*, or *IFIT3* was associated with poor survival in bladder cancer patients, with *IFIT3* also showing a trend toward lower survival despite not reaching statistical significance (Fig. 5G). The remaining seven genes did not show significant associations with patient survival (Fig. S1). Collectively, these analyses suggest that members of the IFIT gene family, particularly *IFIT1*, *IFIT2*, and *IFIT3*, are closely linked to CTS-associated transcriptomic changes and adverse clinical outcomes in bladder cancer, highlighting their potential relevance in CTS-related antitumor mechanisms.

**Fig. 5.**
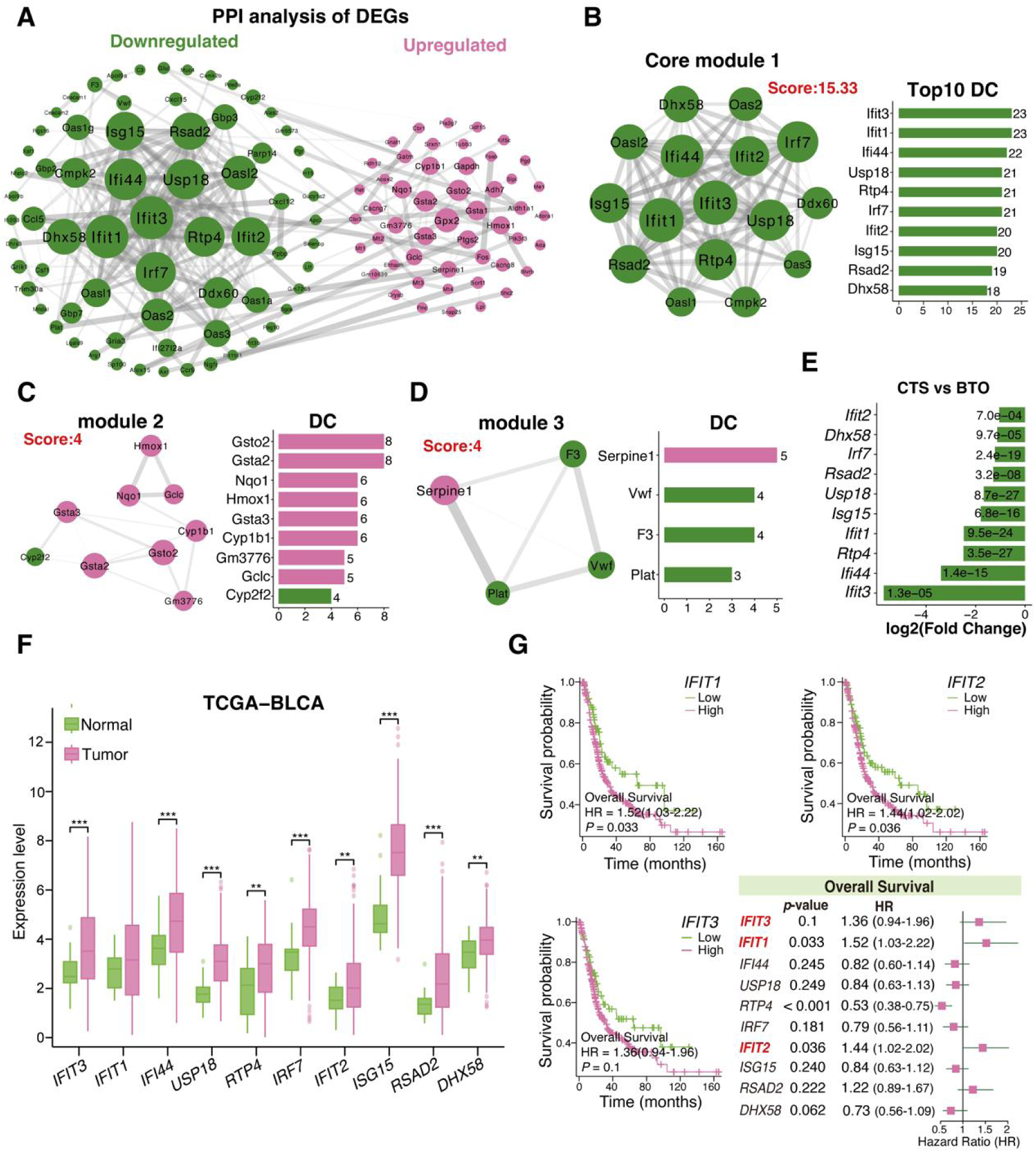
Through DEG-based PPI network analysis, IFIT1/2/3 emerged as central nodes within CTS-associated networks in bladder cancer. **A** PPI analysis of DEGs, comprising 120 nodes and 296 edges, with green representing downregulated genes and red representing upregulated genes. **B** PPI of core module 1 (left) identified by MCODE, with the top10 targets ranked by degree value (right). **C** PPI of module 2 (left) identified by MCODE, with the targets ranked by degree value (right). **D** PPI of module 3 (left) identified by MCODE, with the targets ranked by degree value (right). **E** Log2(Fold Change) and p-values of the top 10 targets in core module 1 between CTS and BTO. **F** mRNA expression of the human counterparts of the top 10 targets in core module 1 in tumor tissues and adjacent normal tissues of bladder cancer (TCGA-BLCA). **G** Kaplan-Meier survival curves comparing high and low expression levels of *IFIT1/2/3* in bladder cancer from the TCGA-BLCA dataset, and a forest plot was used to analyze the human counterparts of the top 10 targets in core module 1 with respect to Overall Survival. * *P* < 0.05, ** *P* < 0.01, *** *P* < 0.001, **** *P* < 0.0001.

### 3.6. IFIT1/2/3 are upregulated in bladder cancer and Correlate with immune infiltration

By systematically analyzing RNA-seq data from multiple cancer types in The Cancer Genome Atlas (TCGA), we evaluated the expression patterns of IFIT1, IFIT2, and IFIT3 in tumor tissues compared with adjacent normal tissues. IFIT1, IFIT2, and IFIT3 were consistently expressed at higher levels in tumor tissues across multiple cancer types, including bladder cancer (Fig. S2A). Based on Cancer Cell Line Encyclopedia (CCLE) data, Based on Cancer Cell Line Encyclopedia (CCLE) data, we further examined the relative expression of IFIT1, IFIT2, and IFIT3 across different cancer cell lines (Fig. S2B), as well as specifically in bladder cancer cell lines (Fig. S2C).

Analysis of TCGA-BLCA transcriptomic and clinical data showed that the expression levels of IFIT1, IFIT2, and IFIT3 were higher in patients with advanced clinical stages compared with those at early stages, and were also elevated in tumors with high histological grade (Fig. S2D) and in the non-papillary subtype (Fig. S2E). These observations indicate that increased IFIT expression is associated with more aggressive clinicopathological features in bladder cancer. Given that IFIT family members are interferon-stimulated genes previously implicated in immune-related processes and cancer biology[56], we further examined the relationship between IFIT expression and immune-related characteristics in bladder cancer. Using the ESTIMATE algorithm[42], the expression levels of IFIT1, IFIT2, and IFIT3 showed significant positive correlations with ESTIMATE score, immune score, and stromal score (Fig. 6A; Fig. S3A), along with a significant negative correlation with tumor purity (Fig. S3B). Consistently, immune-related enrichment scores were higher in the IFIT1-, IFIT2-, or IFIT3-high expression groups than in the corresponding low-expression groups (Fig. 6B). Further analyses demonstrated that the expression of IFIT1, IFIT2, and IFIT3 was positively correlated with the estimated infiltration of multiple tumor-infiltrating immune cell populations, including CD8⁺ T cells, neutrophils, macrophages, regulatory T cells (Tregs), NK cells, and myeloid dendritic cells (all P < 0.001; Fig. 6C, Fig. S3C–Fig. S4). In line with these findings, the abundance of most tumor-infiltrating lymphocyte populations was higher in tumors with high IFIT1, IFIT2, or IFIT3 expression compared with those with low expression (Fig. 6D). In addition, the expression levels of IFIT1, IFIT2, or IFIT3 were positively correlated with a broad range of immune cell marker genes (Fig. 6E, Fig. S5), further supporting their potential role in regulating immune activity within the tumor microenvironment.

**Fig. 6.**
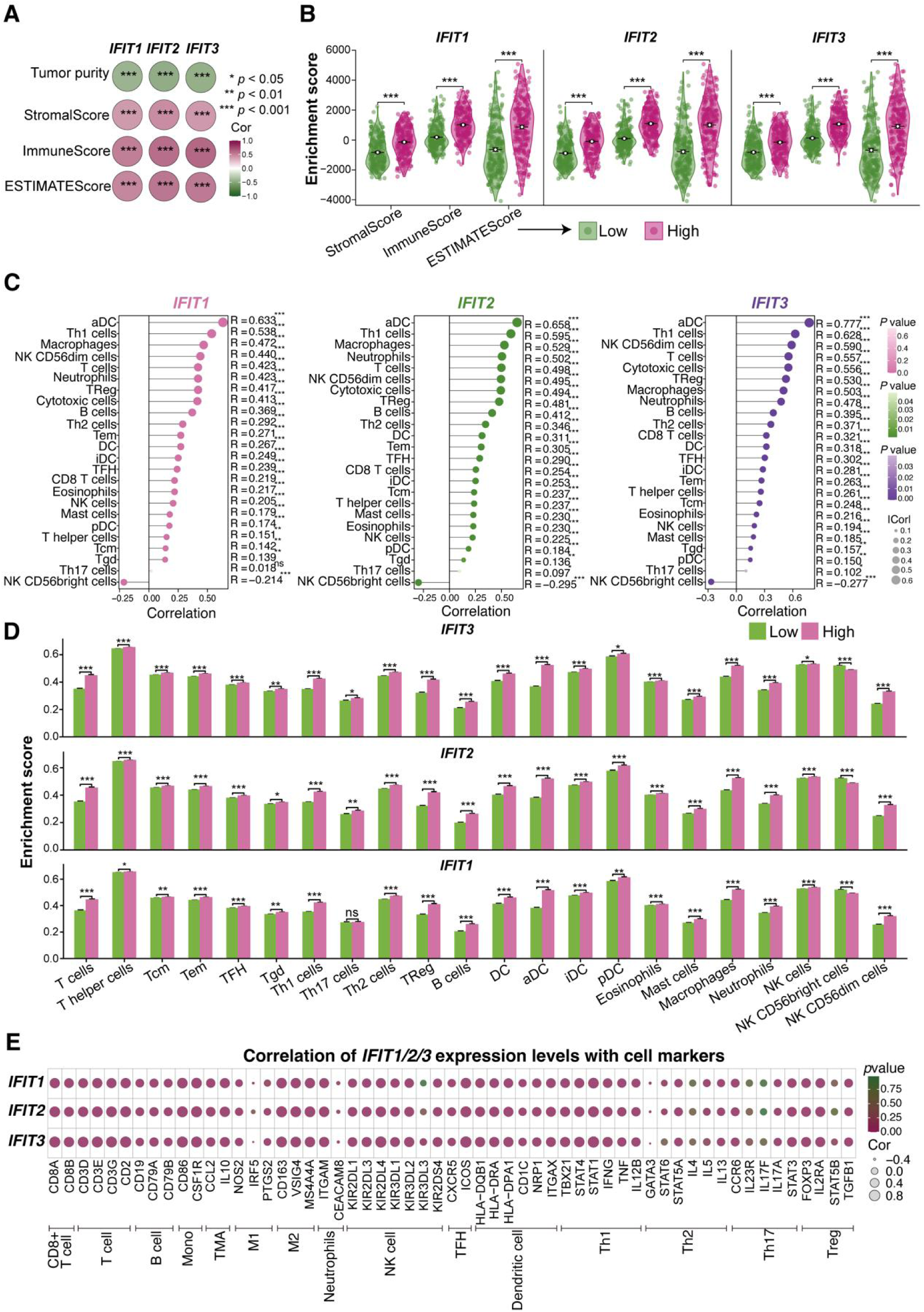
Analysis of the correlation between the expression levels of *IFIT1/2/3* and immune cell infiltration levels in bladder cancer. (A) Correlations between the expression levels of *IFIT1/2/3* and ESTIMATE, Immune, Stromal scores and Tumor purity in BLCA are presented in a heatmap. (B) A violin plot comparing ESTIMATE, Immune, and Stromal scores of immune infiltration between high and low expression groups of IFIT1/2/3 in bladder cancer. (C) The lollipop plot shows the relationship between the abundance of 24 immune cells and *IFIT1/2/3/* expression in bladder cancer, with analysis performed using the ssGSEA immune infiltration algorithm. (D) The bar chart shows the comparison of immune cell infiltration scores in 24 immune cell types between high and low expression groups of IFIT1/2/3 in bladder cancer. (E) Bubble plot showing the correlation between the expression levels of IFIT1/2/3 and immune cell marker expression in bladder cancer. * *P* < 0.05, ** *P* < 0.01, *** *P* < 0.001.

### 3.7. Association between IFIT1/2/3 expression and immune-related factors in bladder cancer

Immunoregulatory factors, chemokines, and major histocompatibility complex (MHC) molecules are key components of the tumor immune microenvironment and are closely related to immune cell recruitment and activation[57, 58]. Chemokines and their receptors coordinate the trafficking and spatial organization of immune cells within tumors[59], whereas MHC molecules are involved in antigen presentation and subsequent T-cell activation, which are essential for antitumor immune responses[58, 60]. In parallel, immune checkpoint and co-stimulatory molecules modulate the balance between immune activation and inhibition, thereby influencing immune surveillance and immune escape[57, 61]. To further examine the immunological associations of IFIT family members, we analyzed the correlations between IFIT1, IFIT2, or IFIT3 expression and a broad range of immune-related factors, including chemokines and their receptors, MHC molecules, and immunostimulatory as well as immunoinhibitory molecules. The expression levels of IFIT1, IFIT2, and IFIT3 showed significant positive correlations with most chemokines (Fig. 7A) and their corresponding receptors (Fig. 7B), such as CXCL10, CXCL11, CXCL4, CCR5, and CCR1. In addition, IFIT1, IFIT2, and IFIT3 expression was positively correlated with multiple MHC-related genes (Fig. 7C), including TAP1, B2M, and HLA-B. Furthermore, IFIT1, IFIT2, and IFIT3 expression levels were positively correlated with markers of immune activation (Fig. 7D), such as TNFSF13B, ICOS, and CD80, as well as with several immune inhibitory molecules (Fig. 7E), including PDCD1, CD274, HAVCR2, and LAG3. Collectively, these findings demonstrate that IFIT expression is broadly correlated with multiple classes of immune-related factors in bladder cancer, reflecting a close association between IFIT family members and immune regulatory features within the tumor microenvironment.

**Fig. 7.**
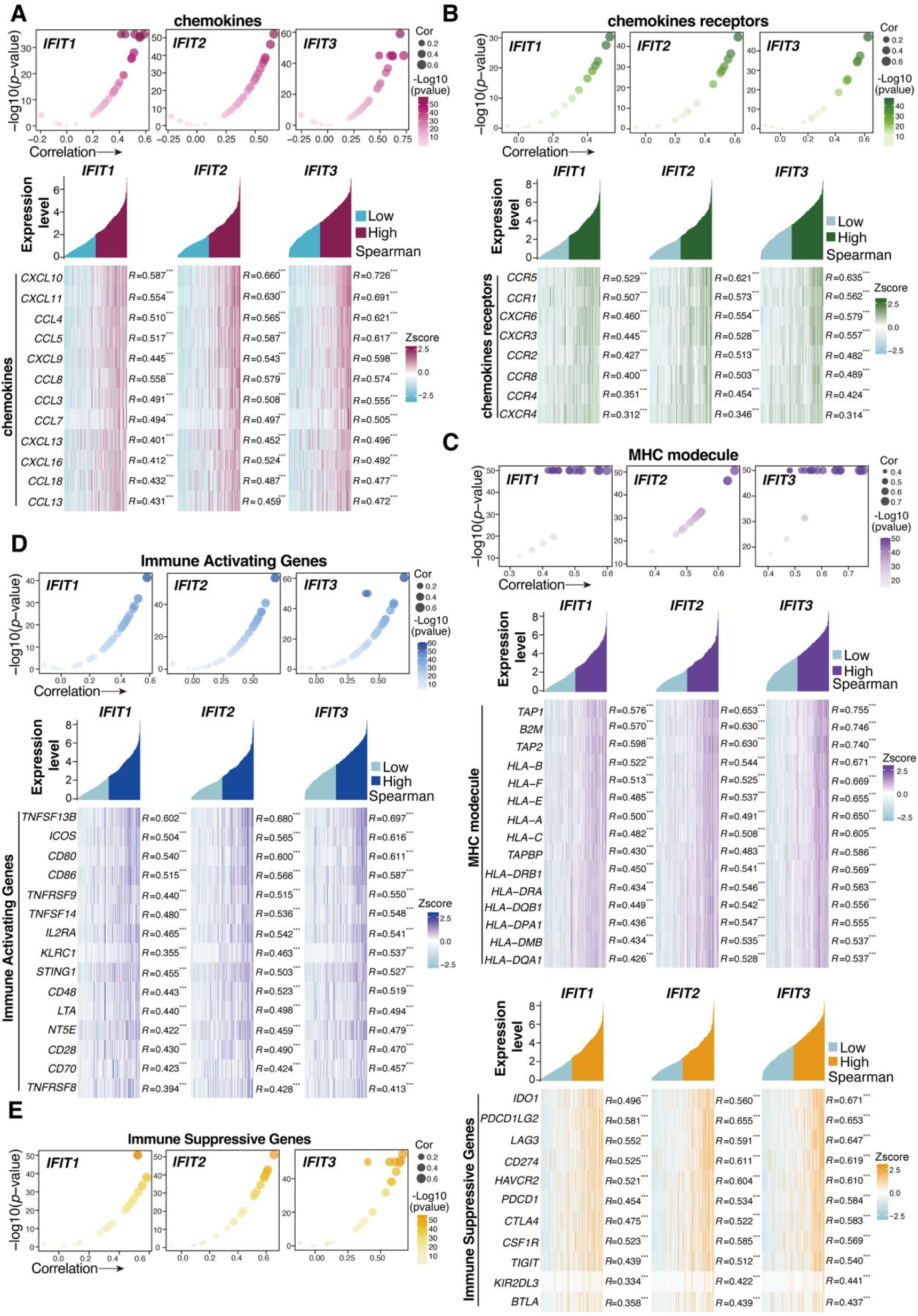
Correlations between the expression levels of *IFIT1/2/3* and immune-related genes. Analyzed genes included chemokines (**A**), chemokine receptors (**B**), MHC modecule (**C**), immune-activating genes (**D**) and immunosuppressive genes (**E**). *P < 0.05, **P < 0.01, ***P < 0.001.

### 3.8. Molecular docking suggests potential interactions between CTS and the cGAS–STING/JAK–STAT signaling axis related to IFIT1/2/3 regulation

To explore molecular mechanisms potentially involved in the regulation of IFIT1/2/3 expression, a GeneMANIA-based analysis[62] was performed to identify genes functionally associated with IFIT1, IFIT2, and IFIT3. This analysis identified 20 genes potentially linked to IFITs, including STING1, STAT1/2, and IRF9 (Fig. 8A). Functional annotation indicated that many of these genes belong to interferon-stimulated genes (ISGs), which are known to participate in antiviral responses and immune-related processes in cancer[63, 64]. Consistently, co-expression analysis using TCGA-BLCA data revealed significant positive correlations between IFIT1/2/3 and these associated genes (R = 0.3–0.8, all P < 0.0001; Fig. S6), suggesting that IFIT family members are embedded within a broader interferon-responsive transcriptional network. Previous studies[65–69] have demonstrated that cyclic GMP–AMP synthase (cGAS) detects aberrant cytoplasmic double-stranded DNA arising from viral infection, pathogenic invasion, or genomic instability in tumor cells, thereby activating the STING1–TBK1–IRF3/7 signaling cascade and inducing type I interferon (IFN-I) production. IFN-I subsequently activates downstream JAK–STAT signaling, leading to the formation of the ISGF3 transcriptional complex (STAT1–STAT2–IRF9), which drives the expression of ISGs, including members of the IFIT family. Together, these established pathways indicate that IFIT expression is closely linked to the cGAS–STING–IFN-I–JAK–STAT signaling axis (Fig. 8B). To gain insight into whether CTS might interact with key components of this regulatory axis, molecular docking analyses were performed using AutoDock Vina[50, 51] to evaluate potential interactions between CTS and representative proteins within the cGAS–STING and JAK–STAT pathways. Binding affinity thresholds (< −4.25 kcal/mol for standard binding, < −5.0 kcal/mol for good binding, and < −7.0 kcal/mol for strong binding) were applied based on established criteria[70]. Within the cGAS–STING pathway, CTS showed predicted binding to STING1 and cGAS. CTS formed three hydrogen bonds with STING1 residues ARG-238 and THR-263, with a binding energy of −8.7 kcal/mol, while docking with cGAS involved four hydrogen bonds at SER-380, GLU-383, and LEU-377, yielding a binding energy of −8.3 kcal/mol (Fig. 8C). Similarly, CTS showed predicted binding to several components of the JAK–STAT pathway, including JAK1, TYK2, and STAT1. Specifically, CTS formed hydrogen bonds with JAK1 residues GLU-883 and SER-963 (binding energy: −9.9 kcal/mol), with TYK2 residues GLN-597 and ARG-738 through five hydrogen bonds (binding energy: −8.5 kcal/mol), and with STAT1 residues ARG-482, THR-451, THR-450, and LYS-240 through five hydrogen bonds (binding energy: −7.3 kcal/mol) (Fig. 8D). Collectively, these docking analyses suggest that CTS has the potential to interact with multiple key proteins within the cGAS–STING and JAK–STAT signaling pathways. These predicted interactions provide a structural basis supporting the possibility that CTS may influence IFIT-associated interferon signaling, thereby contributing to its immunomodulatory effects observed in bladder cancer models.

**Fig. 8.**
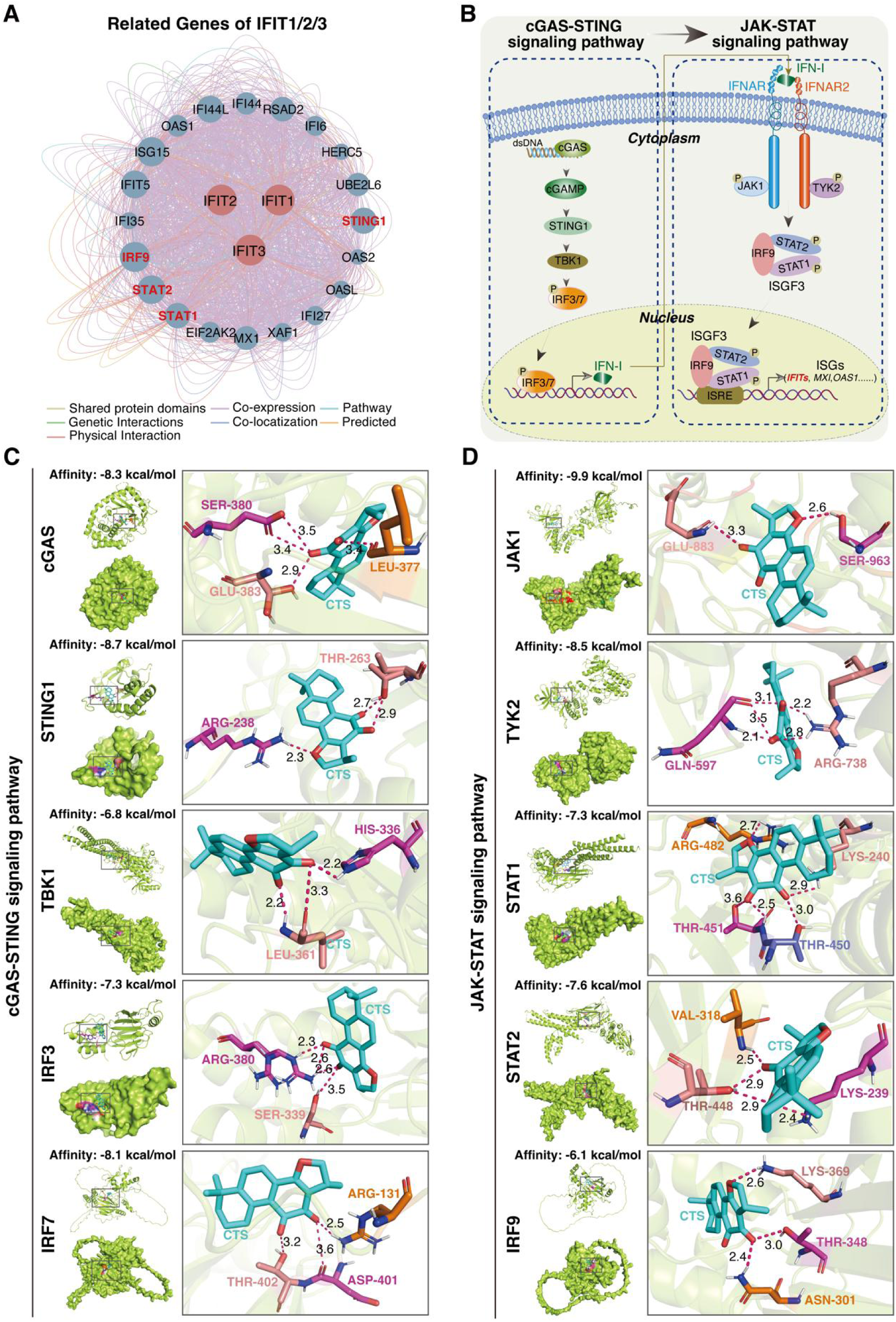
Association between CTS treatment and IFIT family gene expression in relation to cGAS–STING–JAK–STAT signaling pathways. (A) The gene network associated with IFIT1/2/3 analyzed using the GeneMANIA database and visualized through Cytoscape. (B) Diagram of the cGAS-STING-IFN-I-JAK-STAT signaling axis. (C) Molecular docking analysis of key proteins in the cGAS-STING signaling pathway with CTS. (D) Molecular docking analysis of key proteins in the JAK-STAT signaling pathway with CTS.

## 4. Discussion

Bladder cancer (BC) is the ninth most common malignancy worldwide, and the management of muscle-invasive bladder cancer (MIBC) and metastatic disease remains challenging due to difficulties in early detection, high recurrence rates, and limited therapeutic options. Although chemotherapy and radical cystectomy continue to represent standard treatment approaches, their clinical benefits are often constrained by treatment resistance and disease relapse[71]. In recent years, immune checkpoint inhibitors (ICIs), including pembrolizumab[72] and nivolumab[73], have expanded therapeutic options for patients with advanced bladder cancer; however, only a subset of patients derives durable clinical benefit. Consequently, there is a growing need to identify novel therapeutic strategies and biomarkers that may improve patient stratification and treatment outcomes.

During the screening of traditional Chinese medicine monomer compounds for anti-tumor effects, we discovered that cryptotanshinone (CTS) exhibits significant anti-tumor activity. We further investigated its anti-tumor mechanism using a BBN-induced bladder cancer organoid model.

In this study, we identified cryptotanshinone (CTS) as a small-molecule compound with reproducible antitumor activity in bladder cancer models and explored its biological effects using a BBN-induced bladder cancer organoid system. Following initial screening, CTS was selected for further functional evaluation. In conventional two-dimensional bladder cancer cell models, CTS treatment was associated with reduced cell viability in a time- and dose-dependent manner, accompanied by decreased clonogenic capacity, impaired migratory behavior, and increased apoptotic cell populations. These observations indicate that CTS influences multiple cellular processes relevant to tumor growth and survival. Importantly, these effects were also observed in a three-dimensional bladder tumor organoid model, which captures key features relevant to the tumor microenvironment[74]. In this system, CTS treatment was accompanied by reduced organoid growth and decreased expression of the proliferation marker Ki67.

Transcriptomic profiling of bladder cancer organoids revealed that CTS treatment was associated with coordinated changes in gene expression programs related to cell proliferation and immune-associated processes. Notably, genes such as *Ifit1*, *Ifit2*, *Ifit3*, *Cxcl15*, and *Ifi44* showed reduced expression following CTS exposure. These genes have been previously implicated in tumor progression and immune-related signaling pathways in multiple cancer contexts, suggesting that CTS may modulate proliferation- and immune-associated transcriptional programs in bladder cancer cells. Collectively, these findings indicate that CTS elicits consistent transcriptional responses across distinct experimental systems, supporting the rationale for further mechanistic investigation.

The interferon-induced protein with tetratricopeptide repeats (IFIT) family, including IFIT1, IFIT2, IFIT3, and IFIT5, comprises a group of interferon-stimulated genes regulated downstream of interferon signaling pathways and has been implicated in tumor progression, immune regulation, and treatment resistance in several malignancies[24]. In our study, protein–protein interaction (PPI) network analysis highlighted interferon-stimulated genes as prominent components within the CTS-responsive gene network, with IFIT family members occupying relatively central positions. Consistently, the expression of these genes was reduced following CTS treatment in bladder cancer organoids. Analysis of TCGA-BLCA data further revealed that higher expression levels of IFIT family genes were associated with poorer clinical outcomes, supporting their potential relevance to disease aggressiveness.

In addition, immune infiltration analyses demonstrated that the expression levels of IFIT1, IFIT2, and IFIT3 were positively correlated with immune-related scores and inversely correlated with tumor purity, indicating a close association between IFIT expression and immune cell infiltration within the tumor microenvironment. Further immune deconvolution analyses revealed positive correlations between IFIT expression and multiple tumor-infiltrating immune cell populations, including CD8⁺ T cells, macrophages, and regulatory T cells. Moreover, IFIT family expression was positively correlated with the expression of immune checkpoint molecules such as PDCD1, CD274, and CTLA4. Together, these findings suggest that elevated IFIT expression is associated with an immunologically active tumor microenvironment that also exhibits features linked to immune regulation and potential immune suppression. This interpretation is consistent with previous reports demonstrating that modulation of upstream immune-regulatory pathways can influence immune checkpoint expression and tumor immune suppression[75, 76]. Emerging evidence further indicates that epigenetic and transcriptional regulators contribute to interferon-dependent immune programs, highlighting the potential relevance of upstream regulatory mechanisms in the control of IFIT gene expression[77].

At the mechanistic level, GeneMANIA-based network analysis, together with prior literature, supports the involvement of the cGAS–STING[78] and JAK-STAT[79] signaling pathways play important regulatory roles in the expression of IFIT family genes. Molecular docking analysis indicated that CTS may bind to key proteins in these pathways (such as cGAS, STING, TBK1, IRF3/7, and JAK1), inhibiting their activity and thereby downregulating the expression of IFIT genes. This finding provides new molecular targets for the anti-tumor mechanism of CTS and theoretical support for future bladder cancer immunotherapy strategies.

## 5. Conclusion

In summary, this study suggests that CTS is associated with inhibitory effects on bladder cancer cell growth in both two-dimensional cell models and three-dimensional organoids. Transcriptomic and bioinformatic analyses indicate that CTS treatment is accompanied by alterations in immune-related gene expression, including reduced expression of IFIT family members, potentially linked to interferon-responsive signaling pathways such as cGAS–STING and JAK–STAT. These findings are based on computational analyses and organoid models, and further experimental validation in immune-competent in vivo systems is required to confirm the proposed mechanisms and therapeutic relevance of CTS.

## Acknowledgments

We thank TCGA, STRING, TIMER, and TISIDB for offering essential bioinformatics resources. This work was supported by the National Natural Science Foundation of China (Grant No. 82204695).

## CRediT authorship contribution statement

**Mengni Yang:** Data curation, Formal analysis, Investigation, Methodology, Software; Visualization, Writing - original draft. **Rui Li:** Data curation, Visualization, Formal analysis. **Yu Dong:** Validation, Project administration. **Mengting Zhou:** Formal analysis, Validation. **Junning Zhao:** Conceptualization, Methodology, Supervision. **Miao Liu:** Conceptualization, Methodology, Formal analysis. **Ruirong Tan:** Conceptualization, Funding acquisition, Writing - review & editing

## Institutional Review Board Statement

All animal experiments were conducted in accordance with institutional guidelines and approved by the Laboratory Animal Ethics Committee of Sichuan Academy of Chinese Medicine Sciences (Approval No. R20220303-1).

## Data and materials availability

The TCGA-BLCA public data used in this study can be accessed through the UCSC XENA database (http://xena.ucsc.edu/). Data for immune infiltration analysis were obtained from the TIMER database (http://timer.cistrome.org/) and the TISIDB database (http://cis.hku.hk/TISIDB/). The relevant sequencing datasets generated in this study, including transcriptomic data, will be deposited in appropriate public databases for future research use. All data needed to evaluate the conclusions in the paper are present in the paper and/or the Supplementary Materials.

## Informed Consent Statement

Not applicable.

## Abbreviations

The following abbreviations are used in this manuscript

BC: Bladder cancer
CTS: Cryptotanshinone
BCA: Baicalein
BA: Betulinic acid
Tan-IIA: anshinone IIA
PUE: Puerarin
IFITs: interferon-induced tetratricopeptide repeats
BTO: Bladder tumor organoids

