## Supplementary Figures and Supplementary Tables for "Organoid-Based Transcriptomics Indicate IFIT-Associated Immune Modulation during Cryptotanshinone Treatment in Bladder Cancer"

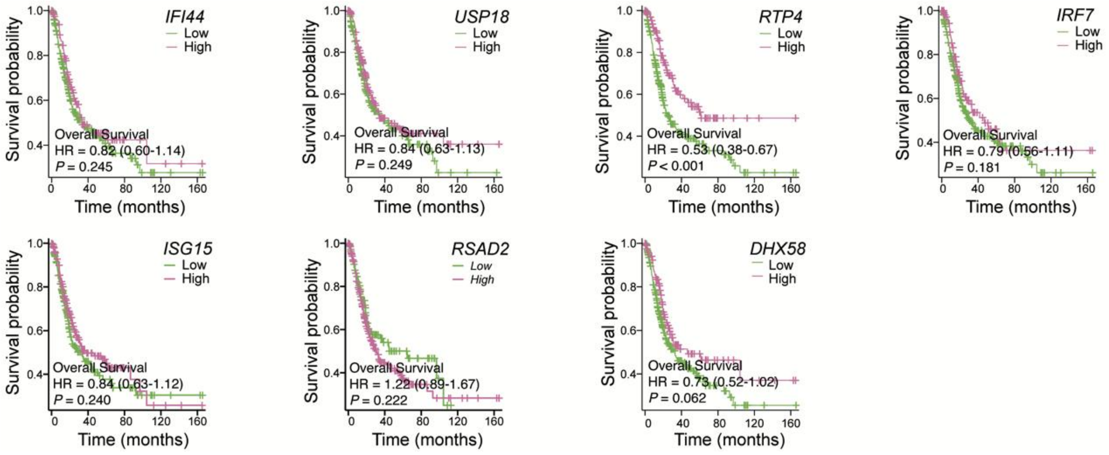


**Fig. S1.** Kaplan-Meier survival curves comparing high and low expression levels of *IFI44*, *USP18*, *RTP4*, *IRF7*, *ISG15*, *RSAD2*, *DHX58* in bladder cancer from the TCGA-BLCA dataset.

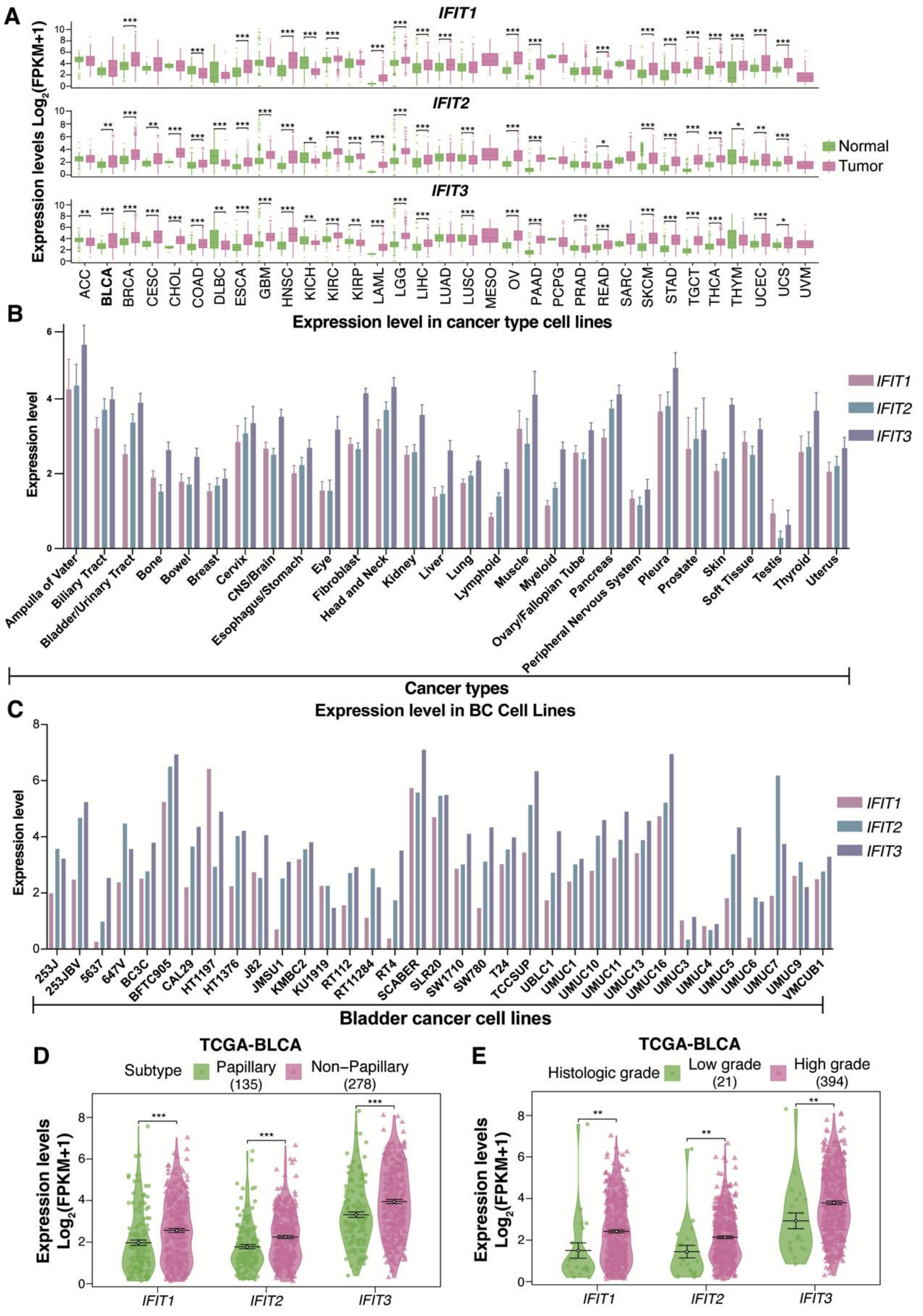


**Fig. S2.** Pan-cancer or Bladder cancer analysis of IFIT1/2/3 expression. (A) The TCGA and GTEx databases were utilized to analyze the expression of IFIT1/2/3 in tumor and normal tissues. (B-C) Expression levels of *IFIT1/2/3* in tumor cell lines from the CCLE database, across different cancer cell types (B), or in various bladder cancer cell lines (C). (D-E) The expression levels of IFIT1/2/3 across subtypes (Papillary: n = 134, Non-Papillary: n = 273) (D) and histologic grades (Low-grade: n = 21, High-grade: n = 388) (E) from TCGA-BLCA. * P < 0.05, ** P < 0.01, *** P < 0.001, **** P < 0.0001.

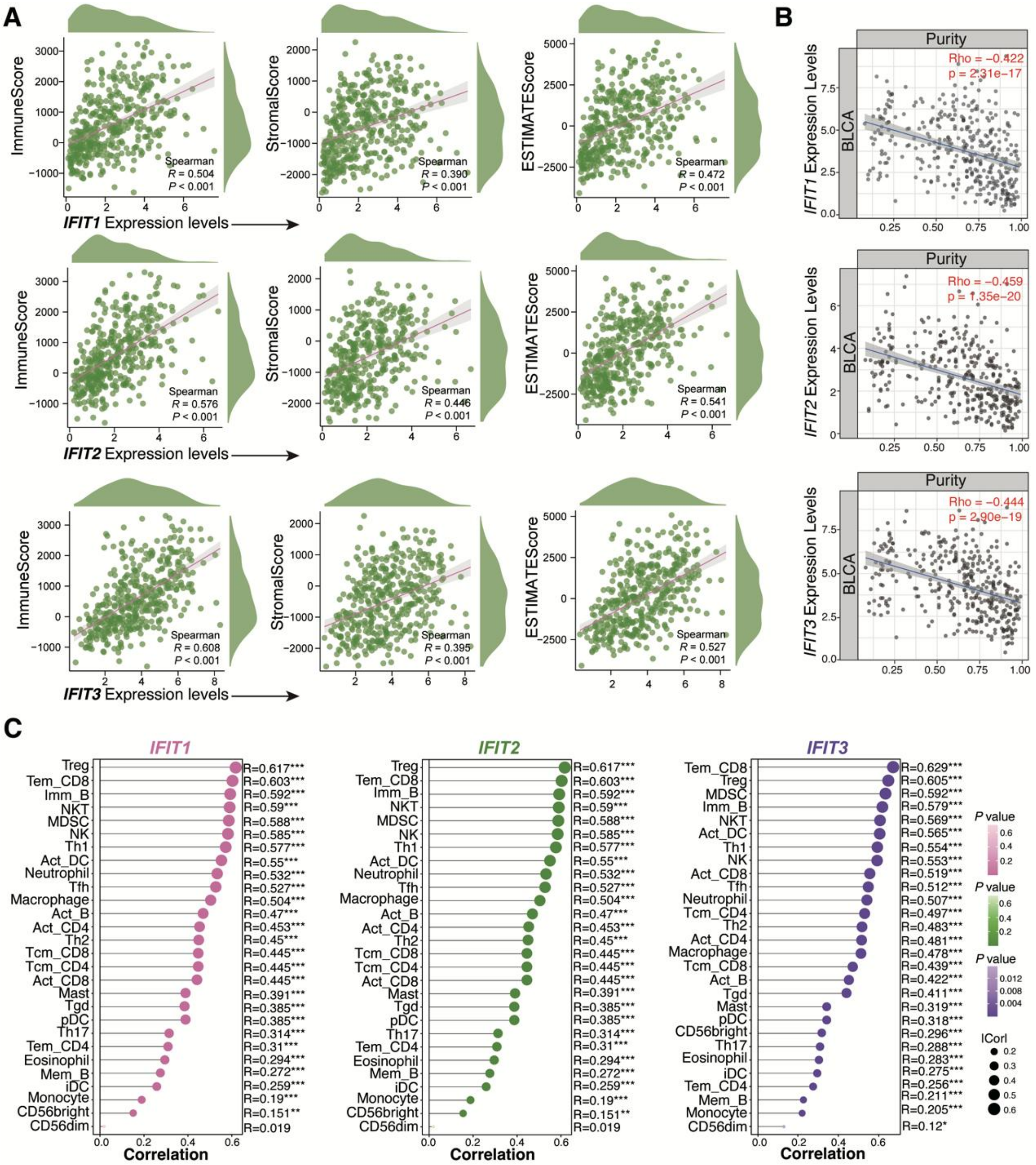


**Fig. S3.** Immune infiltration analysis of IFIT1/2/3 in bladder cancer. (A) Scatter plot of correlations between the expression levels of *IFIT1/2/3* and ESTIMATE, Immune, Stromal scores in bladder cancer. (B) Scatter plot of the correlation between the expression levels of IFIT1/2/3 and tumor purity in bladder cancer. (C) Based on data obtained from the TISIDB database, the lollipop plot shows the relationship between the abundance of 28 immune cells and the expression of IFIT1/2/3 genes in bladder cancer.

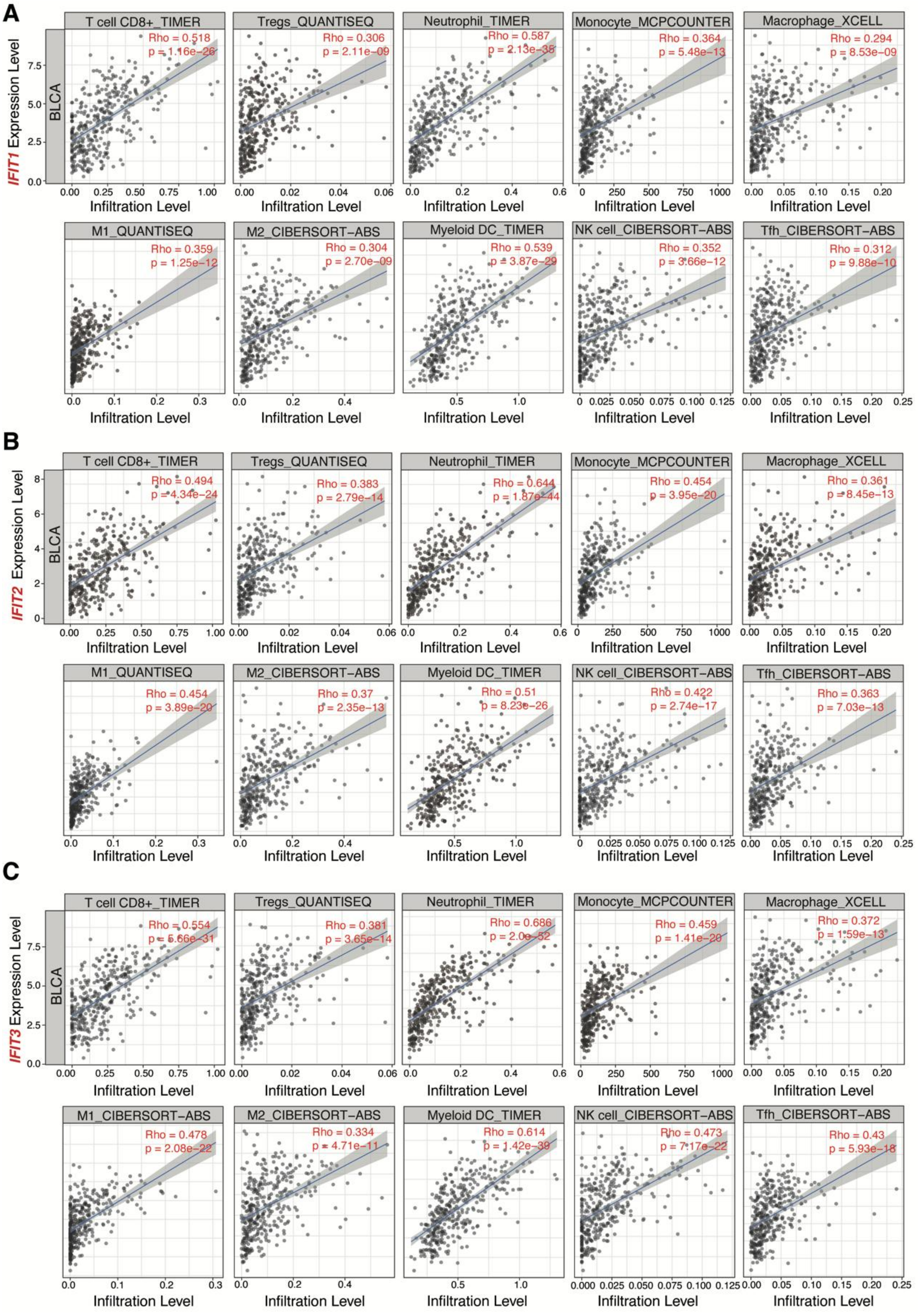

**Fig. S4.** Correlation of IFIT1/2/3 expression with infiltration levels of CD8 + T cell, Treg cell, neutrophil, Monocyte, macrophage, myeloid dendritic cell, natural killer cell in bladder cancer available at TIMER2.0 database.

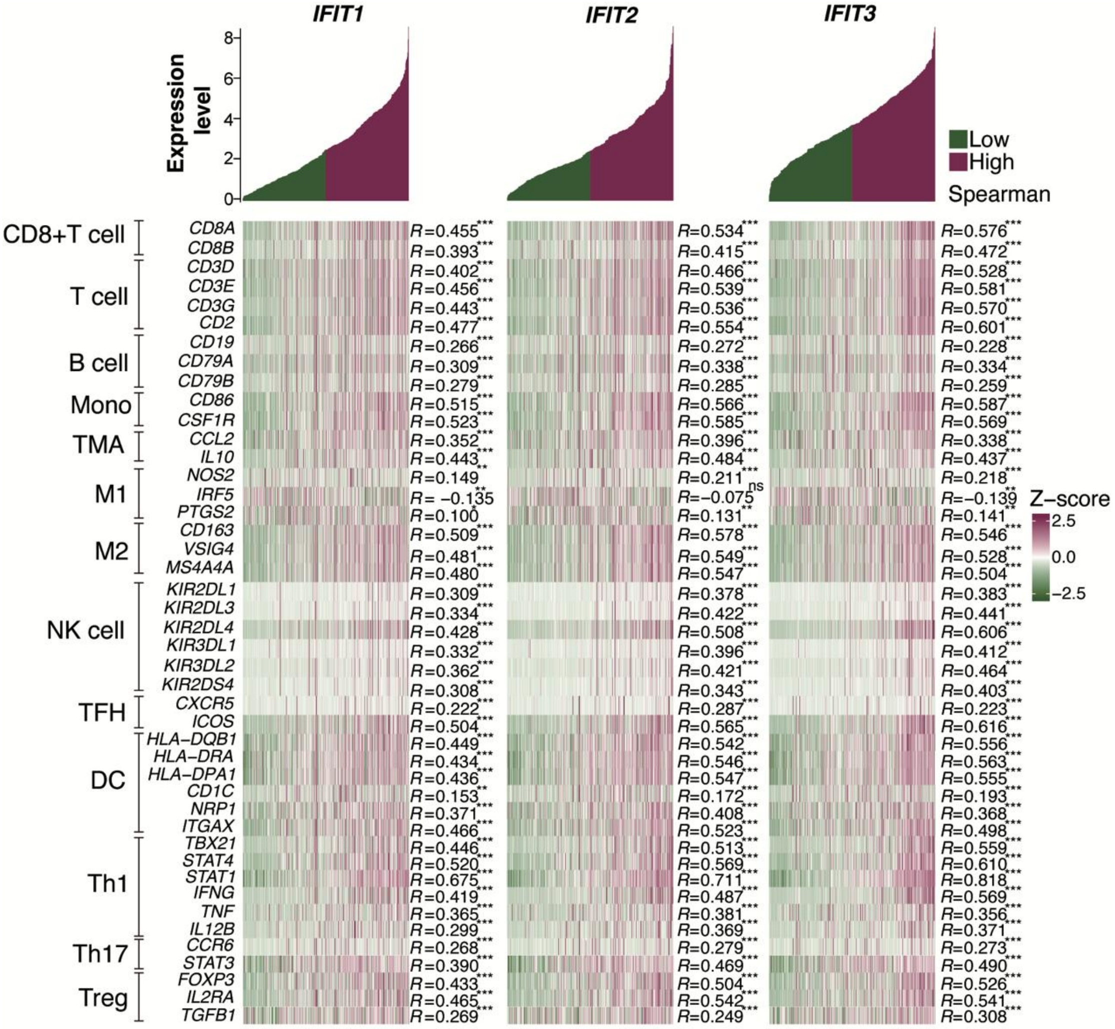


**Fig. S5.** Co-expression analysis of IFIT1/2/3 with immune cell markers, based on Spearman correlation.

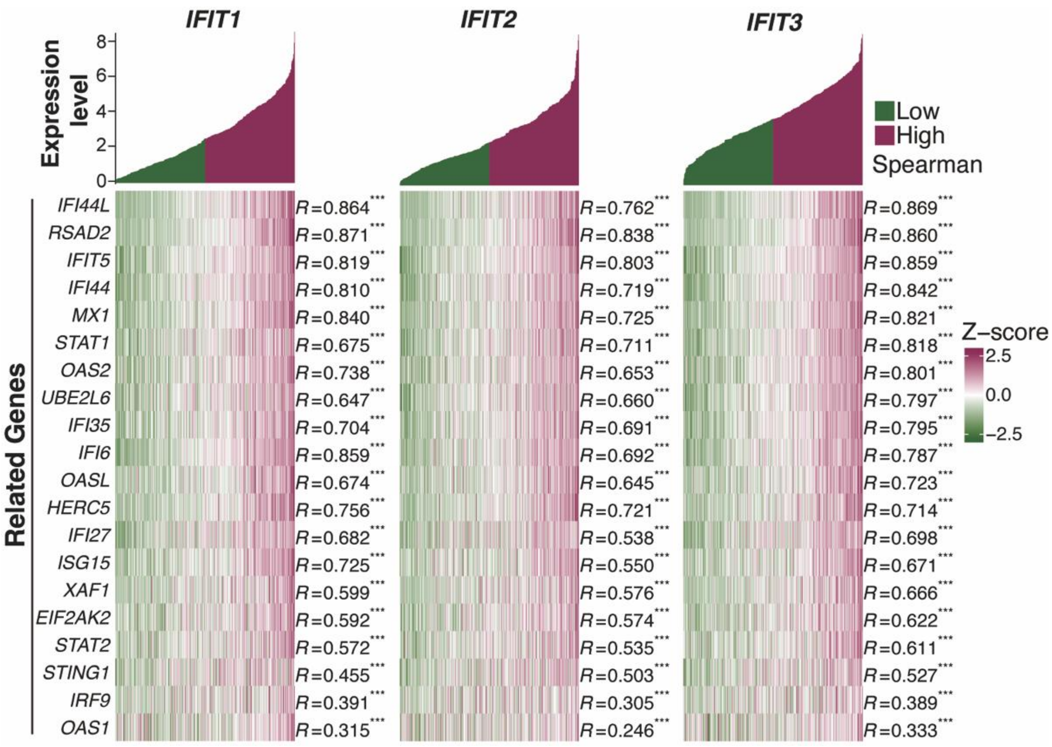


**Fig. S6.** Co-expression analysis of IFIT1/2/3 with related genes, based on Spearman correlation.

### Supplementary Tables

#### Table S1 Target protein pocket coordinates and Grid box sizes

| Target protein | UniProt ID | PDB ID | Protein pocket coordinates | Grid Box size |
|---|---|---|---|---|
| cGAS | Q8N884 | 7FUA | X = 11.853, Y = -10.258, Z = -12.572 | X = 44, Y = 66, Z = 50 |
| STING1 | Q86WV6 | 7KW1 | X = 29.946, Y = 9.404, Z = 12.54 | X = 36, Y = 42, Z = 46 |
| TBK1 | Q9UHD2 | 4IM0 | X = 90.5, Y = 44.631, Z = -16.574 | X = 66, Y = 114, Z = 68 |
| IRF3 | Q14653 | 5JEO | X = -34.258, Y = 5.469, Z = 15.259 | X = 28, Y = 62, Z = 52 |
| IRF7 | Q92985 | AlphaFold | X = -2.027, Y = -1.368, Z = 6.252 | X = 58, Y = 62, Z = 70 |
| JAK1 | P23458 | 6SM8 | X = 9.961, Y = 31.375, Z = 9.697 | X = 40, Y = 80, Z = 40 |
| TYK2 | P29597 | 8TB6 | X = 12.879, Y = 19.483, Z = -30.155 | X = 70, Y = 60, Z = 70 |
| STAT1 | P42224 | 1BF5 | X = 69.706, Y = 36.832, Z = 80.946 | X = 100, Y = 68, Z = 76 |
| STAT2 | P52630 | 6UX2 | X = -30.935, Y = 20.414, Z = 13.69 | X = 100, Y = 70, Z = 76 |
| IRF9 | Q00978 | AlphaFold | X = -2.954, Y = 20.414, Z = 6.516 | X = 68, Y = 70, Z = 76 |

#### Table S2 Top 10 proteins of Core module 1 information

| | Protein names | DC | BC | CC |
|---|---|---|---|---|
| Ifit3 | Interferon-induced protein with tetratricopeptide repeats 3 | 23 | 780.71 | 0.02 |
| Ifit1 | Interferon-induced protein with tetratricopeptide repeats 1 | 23 | 194.10 | 0.02 |
| Ifi44 | Interferon-induced protein 44 | 22 | 54.62 | 0.02 |
| Irf7 | Interferon regulatory factor 7 | 21 | 720.56 | 0.02 |
| Rtp4 | Receptor-transporting protein 4 | 21 | 50.87 | 0.02 |
| Usp18 | Ubl carboxyl-terminal hydrolase 18 | 21 | 37.55 | 0.02 |
| Ifit2 | Interferon-induced protein with tetratricopeptide repeats 2 | 20 | 543.19 | 0.02 |
| Isg15 | Ubiquitin-like protein ISG15 | 20 | 636.74 | 0.02 |
| Rsad2 | Radical S-adenosyl methionine domain-containing protein 2 | 19 | 170.43 | 0.02 |
| Dhx58 | ATP-dependent RNA helicase DHX58 | 18 | 20.43 | 0.02 |
